# Generalisation between motor and declarative memory sequences: A conceptual replication of Mosha & Robertson (2016)

**DOI:** 10.1101/2025.10.15.682521

**Authors:** Sophie Thong, Joshua Hendrikse, Trevor T. -J. Chong, James P. Coxon

## Abstract

Motor and declarative memory systems have been traditionally considered distinct. However, a study by Mosha and Robertson (2016) reported striking evidence of ‘generalisation’ between motor and declarative learning. Specifically, learning improved if the current task (e.g. motor sequence) shared the same high-level ordinal structure as an earlier task (e.g. word list), demonstrating cross-domain transfer of unstable memories. This finding has significant implications for our understanding and conceptualisation of memory taxonomies but has not been replicated. Here, healthy adult participants (*N* = 125) completed a word list and motor sequence task in counterbalanced order with either a shared or distinct sequence structure. In contrast to Mosha & Robertson (2016), we found that a shared ordinal structure between the declarative and motor sequence tasks did not facilitate performance. Overall, our results challenge the robustness of cross-domain generalisation, and underscore the complexity of cross-memory interactions.

## Introduction

The traditional dichotomy is that motor and declarative memories are encoded through distinct cognitive and neural mechanisms (Henke, 2010; Squire, 2004). More recent studies, however, have shown that learning the order of actions and events engage similar neural networks. It can be inferred from behaviour that the encoding of sequential motor and episodic memories, successively or concurrently, evokes interactions between these memories (e.g., Cohen and Robertson (2011); Gasser and Davachi (2023); Mosha and Robertson (2016)). These interactions suggest that ordinal information may be shared between our different memory systems (Robertson, 2022). However, such interactions have been scarcely studied, and extant data provide conflicting evidence on whether information encoded in one memory system can be implicitly transferred to another domain (e.g., Brown and Robertson (2007); Chen et al. (2020); Kamal et al. (2024)).

A prime example of cross-domain interaction is that of generalisation between a motor sequence and a declarative word list (Mosha and Robertson, 2016). Generalisation is the process through which previously learned information is applied to the encoding or learning of a subsequent experience that shares a common feature, thereby enhancing learning or recall of the subsequently encoded memory (Herszage & Censor, 2018; Robertson, 2018). In the key study by Mosha and Robertson, encoding a declarative word list prior to learning an implicit motor sequence led to significantly greater motor skill when both sequences shared an ordinal structure, relative to when the ordinal structure differed between memories. Remarkably, this generalisation effect was also observed when the order of the declarative and motor tasks was reversed, and occurred without participants’ knowledge of the matched structure.

The findings of Mosha and Robertson (2016) suggest a bi-directional generalisation across memory systems. That is, ordinal information can be extracted from one experience and scaffold the encoding of a subsequent memory with a similar structure, even though the elements of the subsequent memory may be qualitatively different (e.g., actions vs. words) (Robertson, 2022). Their findings therefore have significant implications for theoretical frameworks of memory. Importantly, however, this cross-domain generalisation has only been documented once. The primary aim of the present study was to replicate Mosha and Robertson (2016) to corroborate evidence of cross-domain generalisation on the basis of a shared ordinal structure. In line with this seminal study, we hypothesised that participants who learned motor and declarative sequences with a shared ordinal structure would demonstrate better performance and/or recall on the subsequently learned sequence, relative to those who learned these sequences with a different structure.

## Methods

This study was pre-registered on the Open Science Framework on 13 June 2024 (link: [OSF pre-registration](https://osf.io/9tzjg/?view_only=b6c485e6fdc94d1fba0fd228cb5c7fde)). Our pre-registration included an additional aim which is outside of the scope of this manuscript and is not reported herein.

### Participants

One hundred and twenty-five adults (*M*_age_ = 23.65 ± 4.7 years, 32 male) were recruited via convenience sampling from Melbourne, Australia. All participants were right-handed as determined by the Edinburgh Handedness Inventory (Oldfield, 1971), and did not have a current diagnosis of psychiatric or psychological illness. Ethical approval was obtained by the Monash University Human Research Ethics Committee (Project ID: 38080).

### Rationale for sample size

Our sample size was informed by an *a priori* power analysis in G*Power 3.1.9.7. (Faul et al., 2007) modelled from the effect sizes reported in Mosha and Robertson (2016), i.e., a Cohen’s *d* =1.45 for the transfer effect from word list to motor sequence learning, and a Cohen’s *d* = .94 for transfer effect of motor sequence to word list learning. Experimental power was set to 0.95, for two-tailed independent sample *t*-tests, with an alpha of .05. This analysis revealed that a minimum sample of *N* = 14 was required to detect a transfer effect from the declarative to motor domain, and a minimum sample of *N* = 31 to detect transfer from the motor to declarative domain.

### Design

Following the procedure of Mosha and Robertson (2016), each participant completed a word list recall and motor sequence learning task. We employed a mixed-factorial 2 x 2 design with a between-subject factor of ‘Ordinal Structure’ (same vs. different) referring to whether a shared ordinal structure was present between declarative and motor tasks, and a between-subject factor ‘Task Order’ (word list task followed by motor task vs. motor task followed by word list task) denoting which of the two tasks was administered first (Figure 1a). As such, participants were randomly allocated to one of four groups:

**Fig 1.**
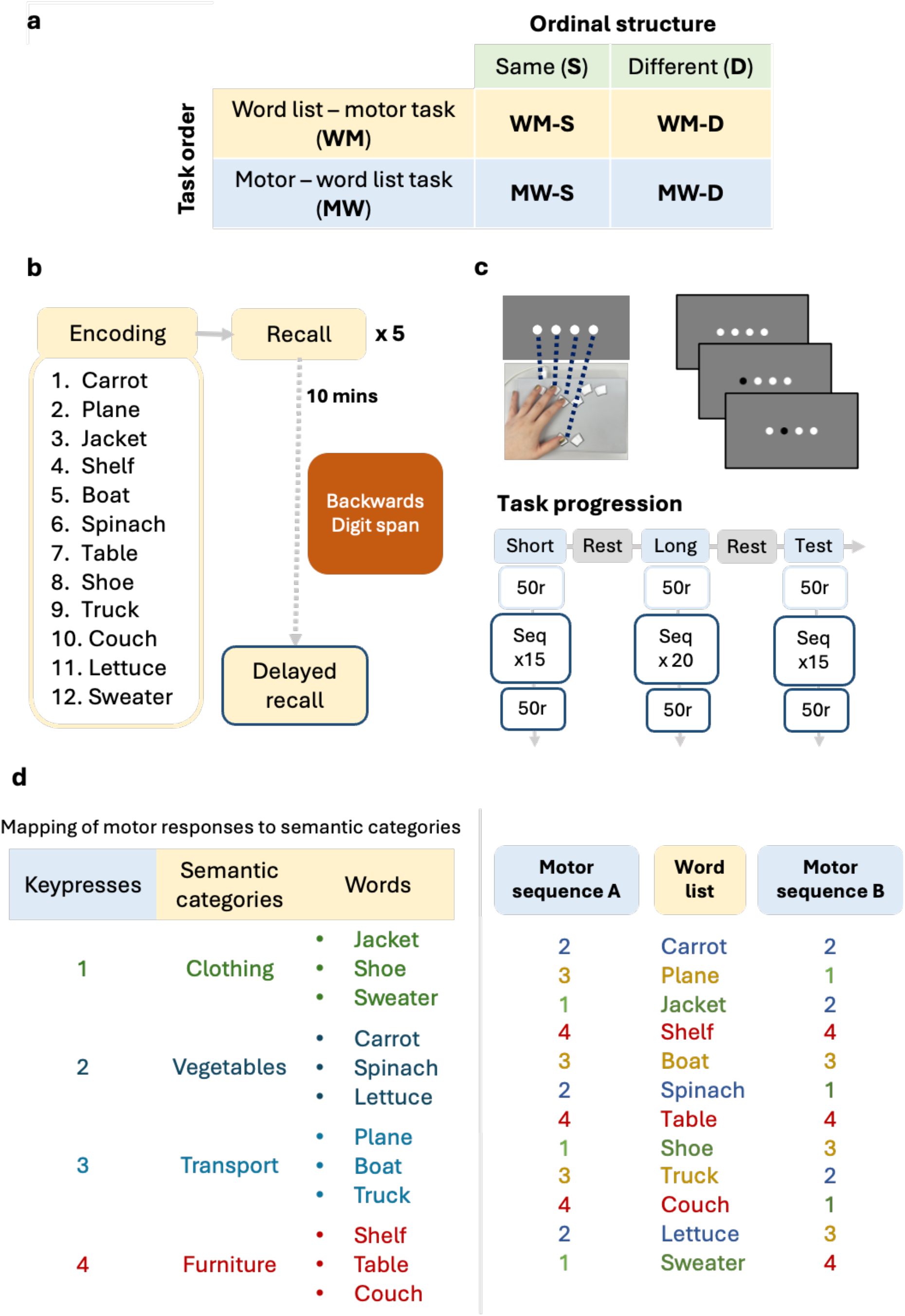
Study methodology. **(a)** Overview of factorial design. Four experimental conditions (groups) were involved which were characterised by Ordinal structure (same (S) or different (D) ordinal structures between the memory tasks) and Task Order (word list task followed by motor task (WM) or motor task followed by word list task (MW)). *W* = word list task, *M* = motor task. **(b)** In the word list task, participants completed five rounds of encoding and immediate recall of the word list. Delayed recall was probed 10 minutes after the fifth recall. During this delay, participants completed the backwards digit span task. **(c)** In the serial reaction time task, participants viewed four circles on a screen which mapped on to buttons on a response box. At the start of each block all circles initially appeared white, followed by a circle turning black to indicate button to be pressed. Participants completed three task blocks, and the motor skill metric obtained on each block was derived from the last 50 pseudorandom and 50 sequential button presses (dark blue outline). **(d)** The mapping of motor sequence keypresses to semantic categories, and their respective exemplar words as per Mutanen et al. (2020). Two motor sequences were presented in the study - one with a shared/matched ordinal structure to the word list (Motor Sequence A) and another which differed (Sequence B). Figure adapted from Mutanen et al. (2020).

- WM-S Group completed the *word list task first* (W) then the *motor task* (M), with both tasks sharing the *Same* (S) ordinal structure (*N* = 31, 9 male)
- WM-D Group completed the *word list task first* (W) then the *motor task* (M), with a *Different* (D) ordinal structure (*N* = 31, 8 male)
- MW-S Group completed the *motor task first* (M) then the *word list task* (W), with both tasks sharing the *Same* (S) ordinal structure (*N* = 31, 8 male)
- MW-D who completed the *motor task first* (M) then the *word list task* (W) with a *Different* (D) ordinal structure (*N* = 32, 7 male)

As the primary aim of this study was to assess the reproducibility of the transfer/generalisation effect between memory tasks/sequences, we did not include a 12-hour re-test session, which was included in the index study (Mosha and Robertson, 2016).

### Procedure

In keeping with the experimental design of Mosha & Robertson (2016), participants completed either a serial reaction time task (SRTT) or word list task first, followed by the other task in succession. All tasks were programmed and administered via PsychoPy (v2023a) (Peirce et al., 2019).

### Word list task

Participants were instructed to remember the list of 12 words (Figure 1b) in the order they were presented on a computer monitor. Each word was displayed for two seconds then replaced by the next word. Following presentation of all 12 words, participants were instructed to recall the word list by typing out as many of the words in their correct order as possible. The word-list presentation and recall were repeated five times. Participants’ recall performance was operationalised as the Levenshtein edit distance, i.e., the number of substitutions, deletions, and insertions required to align participants’ responses with the actual presented word list (Mutanen et al., 2020). A larger edit distance corresponded to poorer recall. For completeness, we also analysed participants’ serial recall, i.e., the longest consecutive sequence of correctly recalled words, as per Mosha and Robertson (2016).

However, we report the edit distance as our main outcome measure on the basis of greater sensitivity/precision relative to serial recall metrics (Gonthier, 2023).

Additionally, as per Mosha and Robertson (2016), we examined participants’ delayed word-list recall 10 minutes after the fifth repetition. In our study, a backwards digit span task was administered to standardise the delay period across participants (in contrast to the index study, for which there was no explicit statement of what happened during the delay). Standardising this delay served to control for variations in participants’ cognitive or brain states during this period (e.g., mind-wandering, active rehearsal of the word list), which may have extraneously influenced their delayed recall (Bradley et al., 2022). During the digit span task, participants heard audio recordings of 16 number lists and verbally reported them in reverse order after each recording. Lists increased in length as the task progressed and all lists were administered to standardise completion time across participant groups. Working memory capacity did not differ significantly between experimental groups (Supplementary Materials).

### Serial reaction time task (SRTT)

In this task, participants were presented with four white circles arranged horizontally on the screen. On each trial, participants were instructed to respond to the circle that turned black by pressing the corresponding button on a response box (Cedrus RB-840, Cedrus Corporation, California) as quickly and accurately as possible. A correct response was required to progress to the next trial (Figure 1c).

Each block of practice started and concluded with 50 trials presented in a pseudo-random order (i.e., randomised while ensuring there were no response repetitions). Undisclosed to participants, the visual cues followed a 12-item repeating sequence through the middle portion of the block (either 2-3-1-4-3-2-4-1-3-4-2-1, or 2-1-2-4-3-1-4-3-2-1-3-4, depending on group allocation as outlined below). Participants completed three blocks of the SRTT (Figure 1c). The first block contained 15 repeats of the sequence, the second was a longer practice block containing 20 repeats, and the final test block comprised 15 repeats. Participants were given a one-minute break between practice blocks. Participants were probed for the development of explicit sequence awareness at the end of the protocol.

Motor sequence learning was operationalised as the difference between participants’ average reaction times (ms) across the last 50 pseudo-random vs. 50 sequential trials. Larger difference values indicate better skill learning (Mosha & Robertson, 2016).

### Matching the ordinal structure of the word list and motor sequence

We adopted the word list from the California Verbal Learning Test as used in Mutanen et al. (2020), which shares the same senior author as Mosha and Robertson (2016). Mutanen et al. (2020) reported the exact word list and keypress sequences (with and without a matched structure) used in their study whereas the original paper did not. The word list included four semantic categories, ‘clothing’, ‘vegetable’, ‘transport’, and ‘furniture’, and these were assigned to keypresses ‘1’, ‘2’, ‘3’, and ‘4’ respectively of the motor sequence task (Figure 1d). As per Mutanen et al. (2020), two motor keypress sequences were utilised -one with a shared ordinal structure between semantic categories and motor responses, and one sequence which differed (Figure 1d). Sequence A had the same ordinal structure, whereas Sequence B had a different ordinal structure. In keeping with Mosha and Robertson (2016) and Mutanen et al. (2020), participants were not informed of the mapping between the two tasks.

### Data analysis

Data pre-processing was conducted using custom MATLAB scripts (version R2022a). Statistical analyses were performed in JASP (v 0.17.2.1) (JASP, 2023).

#### Data pre-processing

For performance on the word list task, participants’ responses were first reviewed and corrected for spelling errors. The Levenshtein edit distance, i.e., the number of substitutions, deletions and insertions required to match participants’ responses to the presented word list (Supplementary Fig 1), was calculated for all responses across all six rounds of recall using custom MATLAB (r2022a) scripts. Additionally, participants’ serial recall, the longest number of words recalled in the correct order was scored manually.

Data from the SRTT were processed according to the approach described in Mosha and Robertson (2016) and Mutanen et al. (2020). Reaction time was defined as the time taken (ms) to respond to each cue after trial onset. For each participant, Grubb’s test was conducted to identify reaction times in the top one percentile, which were removed from subsequent analysis (Mosha and Robertson, 2016, Mutanen et al., 2020). For each practice block, participants’ motor skill (difference between last 50 sequential vs. 50 pseudorandom trials) and accuracy (% correct across last 50 sequential trials) was calculated. Following task completion, participants who were able to correctly report more than five of the 12-item sequence were deemed to have acquired sequence knowledge (Robertson et al., 2004) and were removed from all analyses (see Supplementary Materials).

#### Statistical analysis

To examine transfer from the word list to the SRTT (i.e. declarative to motor), motor skill on the test block was compared between participants in the WM-S and WM-D groups. Conversely, to examine transfer from the SRTT to the word list task (i.e., motor to declarative), we compared delayed word list recall (edit distance and serial recall) between participants in the MW-S and MW-D groups. Independent sample *t*-tests were used for these comparisons, as pre-registered.

In addition to the pre-registered analyses, we also examined whether learning trajectories were influenced by the ordinal structure of the preceding task by comparing learning trajectories when tasks were completed first or following a preceding task. For the SRTT, motor skill and accuracy were the dependent variables. We conducted 2 x 2 x 3 mixed-factorial ANOVAs with the between-subject factors of ‘Task Order’ (motor task first, word task first), ‘SRTT Sequence’ (sequence A, sequence B) and a within-subjects factor of SRTT Block (short, long, and test). For the word list task, separate 3 x 6 mixed ANOVA models were run for the edit distance metric, and the longest number of words recalled in the correct order, with the between-subjects factor ‘Group’ (MW-S Group, MW-D Group, and data pooled across WM-S and WM-D Groups) and the within-subjects factor ‘Recall round’ (Rounds 1 to 6 (delayed recall)).

Mauchly’s test was used to assess the assumption of sphericity and if required, Greenhouse-Geisser corrections (ε < .75) or Huynh-Feldt corrections (ε > .75) were used to adjust the degrees of freedom (Field, 2012). Additionally, Bayes factors (Bayesian independent *t*-test, Cauchy parameter = 0.707) are reported to establish the relative evidence in favour of the null or alternative hypothesis (Ly et al., 2016; Rouder et al., 2009). For significant results, we report BF_10_ values to indicate the strength of evidence for the alternative hypothesis (i.e., that generalisation has occurred), where values ranging from 1 to 3 is considered weak evidence, 3 to 10 moderate evidence and values larger than 10 being strong evidence (van Doorn et al., 2021). For non-significant results, we report BF_01_ values to indicate strength of evidence in favour of the null hypothesis, where values ranging from one to .33 provide weak evidence, .33 to .1 suggest moderate evidence, and values less than .1 indicate strong evidence in favour of the null (van Doorn et al., 2021).

## Results

### Is there evidence of transfer from word to motor sequence learning?

No significant differences in motor skill were observed on the test block of the SRTT when the preceding word list had a matching (WM-S Group: *M* = 72.92, *SD* = 51.26) versus different (WM-D Group: *M* = 58.98, *SD* = 46.92) structure, *t* (56) = -1.078, *p* = .286, *d* = .28, 95%CI = -39.85, 11.97, BF_01_ = 2.31 (Figure 2a). Thus, contrary to Mosha & Robertson (2016), we did not find evidence of facilitated motor sequence learning by virtue of a shared ordinal structure with the preceding word list.

**Fig 2.**
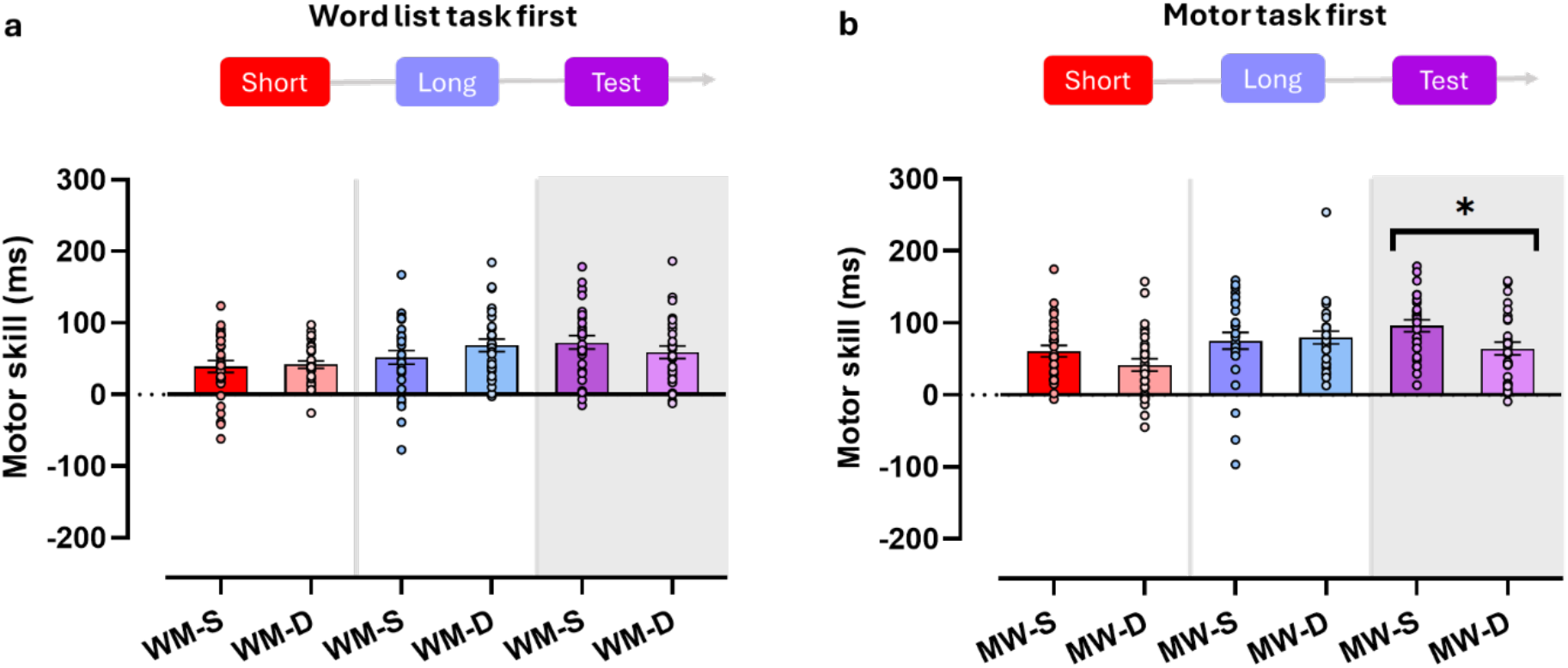
Motor skill for **(a)** those who completed the word list task first and **(b)** those who had completed the SRTT first. ‘Short’, ‘long’ and ‘test’ refer to the three SRTT blocks. Bars and error bars depict *M* ± *SE*. Participants in group WM-S (Panel **a**) completed the same version of the SRTT as participants in group MW-S (Panel **b**). **\*** *p* < .05

Instead, motor sequence learning was influenced by the structure of the motor sequence. Considering just the participants who completed the SRTT first, those who learned Sequence A (MW-S Group skill: *M* = 96.07, *SD* = 42.93) performed better than participants who learned Sequence B (MW-D Group skill; *M* = 64.56, *SD* = 47.79), *t* (55) = -2.615, *p* = .011, *d* = .69, 95%CI = [-55.65, -7.36], BF_10_ = 1.797 (Figure 2b, grey panel). As this result was unexpected, we conducted a post hoc *t*-test to determine if this difference in skill may have been attributable to participants’ attentiveness, approximated by their reaction times on the last 50 pseudorandom trials of the SRTT test block. These groups did not differ significantly in their reaction times (MW-S Group RT: *M* = 548ms, *SD* = 87.7; MW-D Group RT: *M* = 508.2ms, *SD* = 102.51, *p* = .121, BF_01_ = 1.339). Of note, for participants who completed the word list task first and then learned motor Sequence A (WM-S Group), a similar trend was evident (Figure 2a, grey panel). Overall, these findings suggest that motor skill in our sample was largely influenced by motor sequence structure, and not by prior word list learning.

We next examined whether learning trajectories across practice blocks on the SRTT was influenced by the preceding word list task (Figure 2). As expected, all groups improved on the SRTT with practice, *F* (2, 222) = 13.75, *p* < .001, η_p_^2^ = .11. However, the Block x Task Order x Sequence Structure interaction was non-significant, *F* (2, 222) = .093, *p* = .911, η_p_^2^ = 8.39 x 10^-4^, indicating that a shared structure between tasks did not influence the rate of improvement on the SRTT.

Overall accuracy on the SRTT was high, typically >90% (Supplementary Figure 1). There was a slight decrease in accuracy across practice blocks, *F* (2, 222) = 16.07, *p* < .001, η_p_^2^ = .126, and a significant three-way interaction between Block x Task Order x Sequence Structure, *F* (2, 222) = 5.64, *p* = .004, η_p_^2^ = .05. Post-hoc tests comparing groups for each block were not significant (see Supplementary Results).

### Is there evidence of transfer from motor to word sequence learning?

At delayed recall, we observed a trend towards a higher edit distance, i.e., worse performance, when ordinal structure was matched (MW-S Group: *M* = 3, *SD* = 2.38), versus different (MW-D Group: *M* = 1.87, *SD* = 2.1), *t* (56) = -1.97, *p* = .053, *d* = .519, 95%CI = [-2.284, .017], BF_01_ = .756, Figure 3a). Serial recall did not reveal differences between groups (MW-S Group: *M* = 6.36, *SD* = 3.86; MW-D Group: *M* = 7.47, *SD* = 4.01), *t* (56) = 1.07, *p* = .288, *d* = .282, 95%CI = [-.96, 3.18], BF_01_ = 2.32 (Figure 3b). Thus, regardless of how learning was quantified and contrary to expectations (Mosha & Robertson, 2016), we did not observe improved recall when prior motor sequence learning shared the same ordinal structure as the word list.

**Fig 3.**
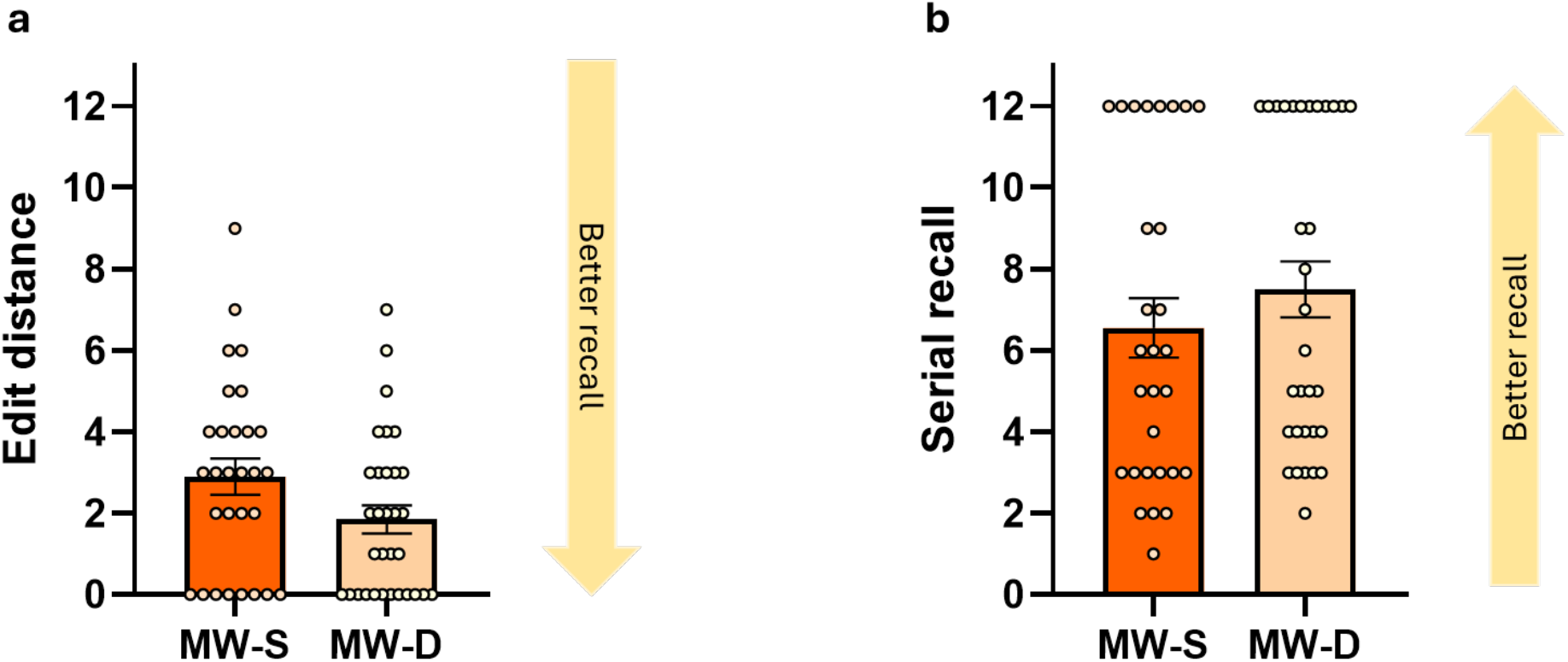
Delayed word list recall of groups MW-S (dark orange bar) and MW-D (light orange bar) operationalised as (**a**) the edit distance and (**b**) serial recall.

To examine whether the trajectory of word list learning was influenced by motor sequence learning, we conducted a mixed-factorial ANOVA with factors Group and Block. For edit distance (Figure 4a) there was a main effect of Block (*F* (3.55, 1903.66) = 191.606, *p* < .001, η_p_^2^ = .62), but no Group x Block interaction (*F* (7.1, 20.36) = 1.024, *p* = .414, η_p_^2^ = .017). A similar pattern of learning was found when serial recall was analysed (Figure 4b), with a main effect of Block (*F* (3.261, 2051.56) = 74.842, *p* < .001, η_p_^2^ = .392), and no Group x Block interaction (*F* (6.521, 67.83) = 1.237, *p* = .284, η_p_^2^ = .02). Altogether, all groups demonstrated similar learning and delayed recall (Fig 4), thus suggesting that the preceding motor structure had a minimal effect on word list learning.

**Fig 4.**
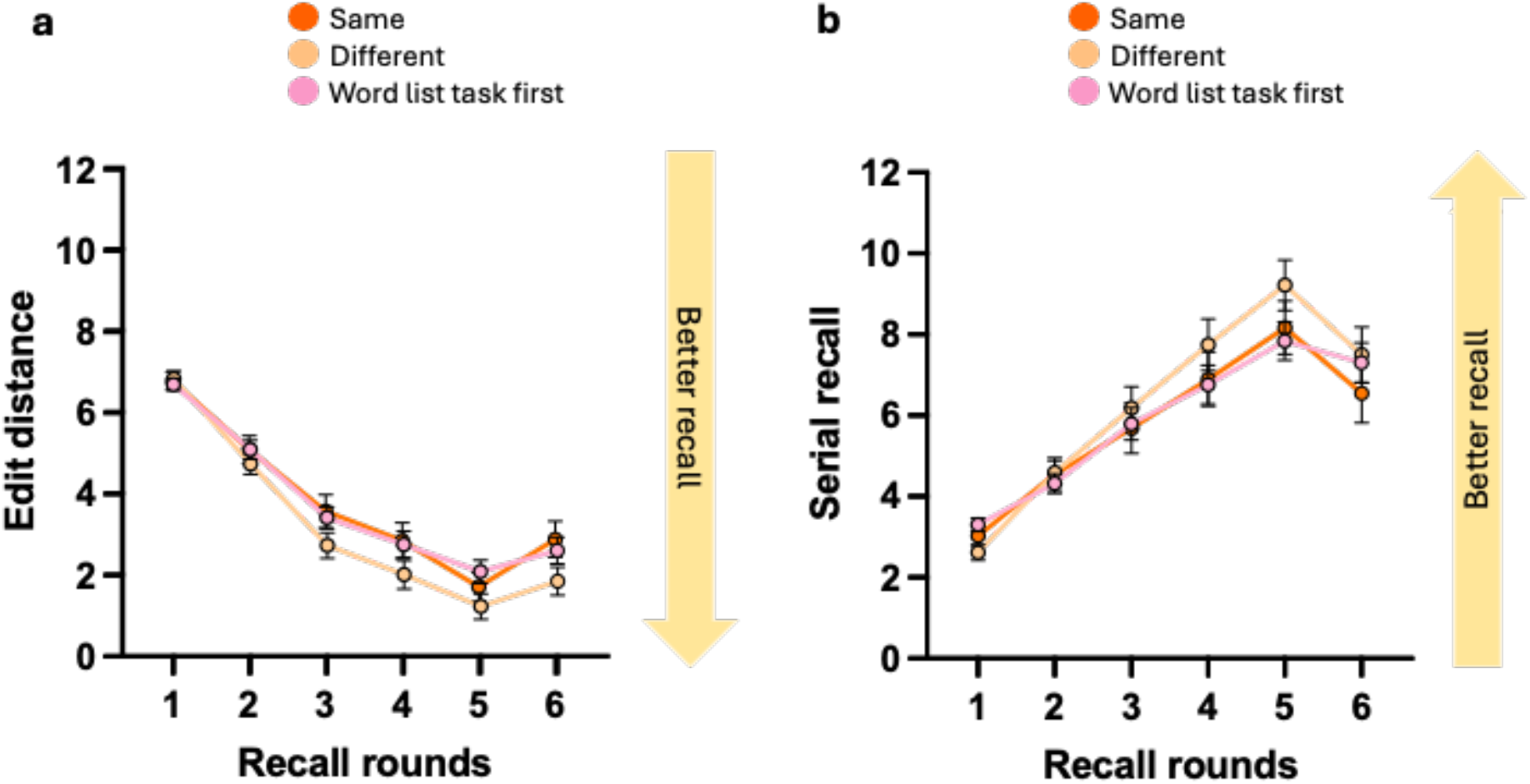
Improvements in word list recall across five successive rounds of encoding and immediate recall for participants who first completed a motor sequence with the same structure as the word list (‘Same’, dark orange line), with a different structure from the word list (‘Different’, light orange line), and participants who completed the word list task first (‘Word list task first’, pink line). (**a)** Recall operationalised as the edit distance, and (**b)** recall operationalised as serial recall.

## Discussion

This study aimed to replicate the findings of Mosha and Robertson (2016) who documented a striking effect of bi-directional generalisation between motor and declarative memory sequences. Contrary to expectations, there was no evidence that a shared ordinal structure facilitated motor or declarative memory formation. The methodology closely followed the index study and was sufficiently powered to detect cross-generalisation effects. Thus, it is unclear why the results diverged from the original study. Regardless, the present results demonstrate that cross-domain generalisation is not a robust effect. The results are discussed in relation to evidence-based theories of hippocampal function and memory interactions.

The absence of generalisation between sequences challenges the idea that overlapping neural ensembles are involved in encoding the ordinal structure of different memory sequences (Robertson, 2022). Cross-domain generalisation is hypothesised to occur when hippocampal ensembles encode the structure of the first memory, and, while still unstable and excitable, become allocated to processing the same structure encountered in a subsequent memory (Robertson, 2022). Learning the order of declarative / episodic memories is known to engage hippocampal networks (Davachi & DuBrow, 2015; Hsieh et al., 2014; Ranganath & Hsieh, 2016) and higher-order sequence learning (e.g., where item *n* is predicted by two preceding items, as per the version of the SRTT we administered), is deemed to be hippocampal-dependent (Curran, 1997; Robertson, 2007; Schendan et al., 2003). However, learning on the SRTT may not be strictly driven by hippocampal ensembles and could involve different cognitive mechanisms (e.g., perceptual learning of stimuli, associations between stimuli and responses) and the development of corresponding sequence representations (Abrahamse et al., 2010) in other brain regions (Gheysen & Fias, 2012; Robertson, 2007). Given the behavioural nature of this research, the exact neural mechanisms engaged during SRTT learning in the current and original sample cannot be known, or if they may have differed between samples. Nonetheless, by virtue of individual differences in learning mechanisms on the SRTT, there may have been differences in our sample in the neural ensembles that were recruited during SRTT learning and word list learning.

Cross-memory generalisation has been documented in other studies (Gasser & Davachi, 2023; Mutanen et al., 2020) and related work has found interference between declarative and motor sequences (Brown & Robertson, 2007; Cohen & Robertson, 2011) (but see Kamal et al. (2024)). As with generalisation, interference is also proposed to occur via overlapping neural ensembles (Herszage & Censor, 2018; Robertson, 2018). Thus, it is worthwhile considering the present study’s findings from the perspective that memory systems do overlap to some degree. Contemporary views within declarative and motor domains posit that encoding two separate memories with a common feature results in memories becoming either integrated or differentiated (King et al., 2019; Schlichting & Frankland, 2017; Schlichting & Preston, 2015). Memory integration is generally considered to evoke generalisation (Schlichting & Preston, 2015), while differentiation prevents interactions between memories (Chanales et al., 2019; Chanales et al., 2017). When both sequences shared a structure, we observed no effect of prior learning on subsequent learning. Thus, it is possible that overlapping sequences became differentiated in our study. Future experimental studies, coupled with neuroimaging techniques capable of assessing similarity of neural memory representations (e.g. Chanales et al. (2019)), may be informative.

One possible distinction between the present study and that of Mosha and Robertson (2016) is what participants did during the 10-minute break to assess delayed word list recall. In the present study, the backwards digit span task was administered to standardise participants’ cognitive state during the delay period. The digit span task may have taxed similar brain regions / resources to those involved in stabilising (consolidating) memory of the word list (e.g., the hippocampus (Davachi & DuBrow, 2015; Genzel & Robertson, 2015)). However, in contrast to the digit span, the word list task required repeated memory retrieval which has been proposed to evoke rapid memory consolidation in cortical hubs (Hebscher et al., 2019). Indeed, average delayed serial recall in the present study was comparable to what Mosha and Robertson (2016) report, suggesting that inclusion of the digit span task did not interfere with participants’ ability to memorise the word list. Regardless of the impact of standardising the delay with the digit span task, the present findings suggest that cross-domain generalisation effects are not robust to methodological differences, and that these effects, should they exist, are subtle.

In conclusion, we investigated the reproducibility of cross-domain memory generalisation, as originally described in Mosha and Robertson (2016). Contrary to expectations, there was no evidence of transfer between memory systems. Our findings highlight that cross-memory interactions are more nuanced than currently understood, and that further research is required to characterise the interactions between our different memory systems.

