## Supplemental file for "Generalisation between motor and declarative memory sequences: A conceptual replication of Mosha & Robertson (2016)"

### Example of edit distance calculation

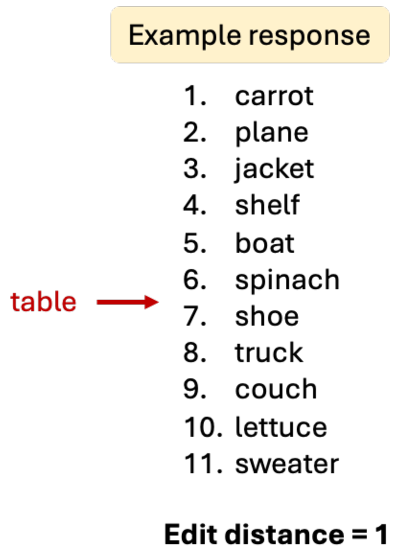


**Supplementary Fig 1.** A participant's word list response requiring the insertion of one word to align it with the original word list and thus results in an edit distance value of 1.

### Working memory (Backwards Digit Span) data

Participants' backwards digit span was scored as the longest list of numbers they accurately repeated in the reverse order. To determine if there may have been extreme differences in working memory across groups, we examined digit span scores using a one-way ANOVA with the Factor: Group. Due to experimenter errors, digit span scores from 10 participants were lost ($N_{Group1} = 2$, $N_{Group2} = 2$, $N_{Group3} = 4$, $N_{Group4} = 2$). No significant differences in digit span performance between groups was observed ($M_{Group1} = 8.62$, $SD_{Group1} = 2.32$; $M_{Group2} = 8.7$, $SD_{Group2} = 2.88$; $M_{Group3} = 8.10$, $SD_{Group3} = 2.09$; $M_{Group4} = 8.63$, $SD_{Group4} = 2.34$), $F\ (3, 111) = .37$, $p = .773$, $\eta^2 = .01$.

### Task exclusions

Participants were excluded from analyses if: 1) they had previously completed the same SRTT or word list task before, 2) they gained explicit knowledge of the motor sequence (able to recall > 5 items of the motor sequence) (Robertson et al., 2004), 3) were detected as outliers by visual inspection of boxplots and/ or 4) for tests of transfer effects, were unable to complete the preceding task correctly (e.g., for analyses examining declarative to motor transfer, failure to complete the word list task correctly resulted in exclusion from SRTT analyses, and vice versa for analyses examining motor to declarative transfer). Exclusions per task by group are reported in Supplementary Table 2.

**Supplementary Table 2**

*Participant exclusions per analysis*

| Participant groups | Tasks | |
|---|---|---|
| | **SRTT** | **Word list** |
| **WM-S** | 1 | 1 |
| **WM-D** | 3 | 0 |
| **MW-S** | 1 | 3 |
| **MW-D** | 1 | 2 |

**Motor sequence accuracy**

Between participant groups WM-S and WM-D, and MW-S and MW-D, post-hoc tests revealed no significant differences in accuracy across the last 50 sequential SRTT trials at any task block (multiple comparisons α = .017, all *ps* > .03).

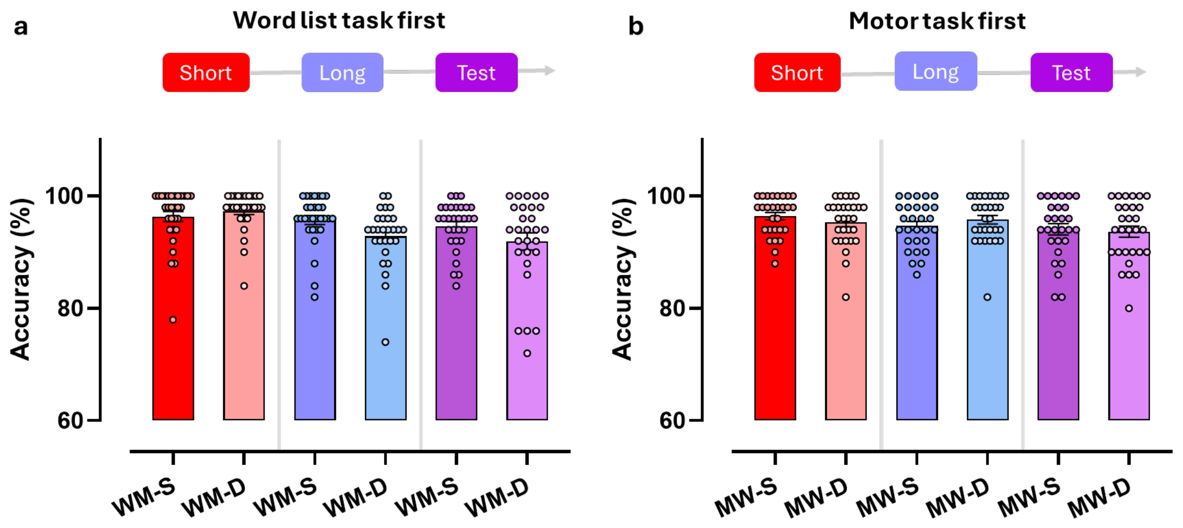

**Supplementary Fig 2.** Accuracy (derived from last 50 sequential trials) on the SRTT for **(a)** participants who completed the word list task first and **(b)** those who had completed the SRTT first. 'Short', 'long' and 'test' refer to the three SRTT blocks. Bars and error bars depict *M* ± *SE.* Participants in the 'same' condition (panel **a**) completed the same version of the SRTT as participants in the 'Sequence A' condition (panel **b**).
